# Interlaboratory Data Variability Contributes to the Differential Principal Components of Human Primed and Naïve-like Pluripotent States in Multivariate Meta-Analysis

**DOI:** 10.1101/822163

**Authors:** Kory R. Johnson, Barbara S. Mallon, Yang C. Fann, Kevin G. Chen

## Abstract

Currently, genome-wide data analyses have revealed significant differences between various human naïve-like pluripotent states derived from different laboratory protocols, confounding the establishment of defining criteria of human naïve pluripotency. Thus, it is imperative to understand the concept concerning the ground or naïve pluripotent state of pluripotent stem cells, which was initially established in mouse embryonic stem cells (mESCs). Putative human pluripotency has been proposed, largely based on comparing genome-wide transcriptomic signatures of human pluripotent stem cells (hPSCs) with human pre-implantation embryos and mESCs by several research groups. Current bioinformatics approaches, however, have inevitable conceptual biases and technological limitations, including the choices of datasets, analytic methods, and interlaboratory data variability. In this report, we performed a multivariate meta-analysis of major hPSC datasets via the combined analytic powers of percentile normalization, principal component analysis (PCA), *t*-distributed stochastic neighbor embedding (*t*-SNE), and SC3 consensus clustering. This vigorous bioinformatics approach has significantly improved the predictive values of the current meta-analysis. Accordingly, we were able to reveal various fundamental inconsistencies between naïve-like hPSCs and their human and mouse *in vitro* counterparts, which are likely attributed to interlaboratory protocol differences. Moreover, our meta-analysis failed to provide global transcriptomic markers that support the putative *in vitro* human naïve pluripotent state, rather suggesting the existence of altered pluripotent states under current naïve-like hPSC growth protocols.

## INTRODUCTION

Ground or naïve states in pluripotent stem cells were initially proposed by Smith and colleagues based on the identification of lineage-primed epiblast stem cells (EpiSCs) derived from post-implantation mouse embryos (Brons et al., 2007; Tesar et al., 2007) and on the studies of naive mouse embryonic stem cells (mESCs) derived from pre-implantation mouse embryos (Nichols and Smith, 2009; Ying et al., 2008). Hence, mouse pluripotent stem cells have two distinct (primed and naïve) states. The maintenance of the naïve state relies on the use of leukemia inhibitory factor (LIF) with two inhibitors, GSK-3βi and ERK1/2i (abbreviated as 2i), that suppress glycogen synthase kinase-3β (GSK-3β) and extracellular signal-regulated kinases 1/2 (ERK1/2), respectively. Conceivably, pluripotent stem cells in the naïve state may have several potential advantages over the primed state, particularly for facilitating single-cell growth, genetic manipulation, disease-modeling, and drug discovery [reviewed in (Chen et al., 2014; Hanna et al., 2010)].

In the past six years, several groups have reported the conversion of primed human pluripotent stem cells (hPSCs), which depend on distinct growth signals that embrace FGF2/Activin-A/TGFβ signaling pathways, to naïve-like hPSCs (NLPs) and *de novo* derivation of NLPs from the human inner cell mass (Chan et al., 2013; Gafni et al., 2013; Guo et al., 2016; Takashima et al., 2014; Theunissen et al., 2016; Valamehr et al., 2014; Ware et al., 2014). However, there is a lack of robust assays that precisely define a naïve pluripotent state under different growth conditions *in vitro*. The existing assays used for defining pluripotent and differentiation states largely count on various genome-wide analyses (Bock et al., 2011; Chan et al., 2013; Gafni et al., 2013; Takashima et al., 2014; Theunissen et al., 2016). However, genome-wide transcriptomic levels across datasets generated from different laboratories using different technologies (e.g., microarray and RNA-sequencing) often have substantial differences in expression scale and spread. Direct meta-analysis of the transcriptomic levels across datasets can render confusing results and lead to incorrect interpretations and conclusions. Thus, previous genome-wide data analyses revealed significant differences between various NLPs derived from different laboratory protocols, thus confounding the definition of human naïve pluripotency (Takashima et al., 2014; Theunissen et al., 2016).

Thus, there is a pressing need to address the above critical issues. In this study, we employed a meta-analysis approach that integrates genome-wide microarrays and RNA sequencing (RNA-seq) data into the principal component analysis (PCA) (Jolliffe, 2002), *t*-student distributed neighboring embedding (*t*-SNE) (van der Maaten and Hinton, 2008), and SC3 consensus clustering (Kiselev et al., 2017). We aim to resolve critical interlaboratory experimental inconsistencies with respect to human naïve pluripotency. With this approach, we characterized transcriptomic signatures of NLPs from publicly available datasets and systematically evaluated data from current human naïve-like protocols. Our analysis revealed the existence of various pluripotent states in converted and derived NLPs, which are deficient of global transcriptomic signatures of naïve pluripotency as described in mESCs and early human embryos. Our study also provides new insights into the role of 1D- and 2D-meta-analysis in gene expression cluster rearrangements, thereby helping define accurate pluripotent states.

## MATERIALS AND METHODS

### Datasets for meta-analysis

We collected 12 datasets for multivariate meta-analysis, including the one (GSE32923) generated by our group (Chan et al., 2013; Gafni et al., 2013; Guo et al., 2016; Mallon et al., 2013; Takashima et al., 2014; Tesar et al., 2007; Theunissen et al., 2016; Valamehr et al., 2014; Vassena et al., 2011; Yan et al., 2013). These datasets are composed of 422 samples from 9 independent laboratories (Supplemental Table 1). The datasets can be identified with GSE and EMBL-EBI accession numbers in parentheses: D1 (GSE32923), D3 (E-MTAB-2031), D5 (GSE46872), D6 (GSE50868), D7 (GSE59435), D18 (GSE7866), D22A (E-MTAB-2857), D22B (E-MTAB-2857), D23 (E-MTAB-2856), D24 (E-MTAB-4461), D25 (GSE36552), and D26 (GSE29397). We further classified these datasets based on their cell types (e.g., blastocysts and human ESCs or hESCs) and cellular states (i.e., primed or naïve pluripotent states) as illustrated in detail in Figure 1.

**Figure 1.**
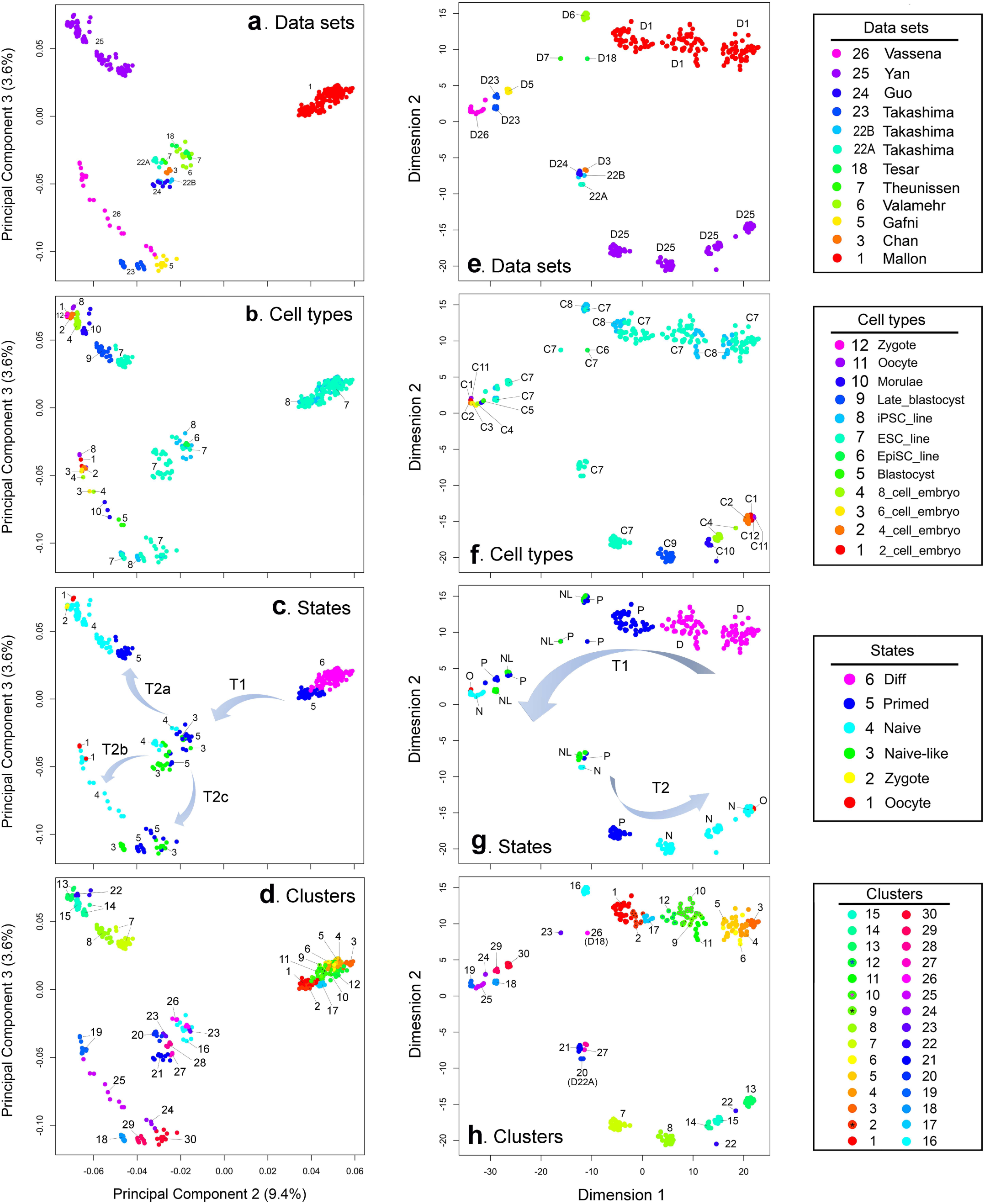
Meta-analysis combining PCA, *t*-SNE, and SC3 consensus clustering for accurately defining pluripotent and cellular states. (The 12 datasets, used in this study, were named based on the first authors of the published reports. The datasets are composed of 422 samples from 9 independent laboratories, which can be identified with GSE and EMBL-EBI accession numbers in parentheses: D1 (GSE32923), D3 (E-MTAB-2031), D5 (GSE46872), D6 (GSE50868), D7 (GSE59435), D18 (GSE7866), D22A (E-MTAB-2857), D22B (E-MTAB-2857), D23 (E-MTAB-2856), D24 (E-MTAB-4461), D25 (GSE36552), and D26 (GSE29397). (**a-d**) PCA and (**e-h**) *t*-SNE visualization plots of all datasets: (right panel) color keys to datasets, cell types, pluripotent and differentiation states, and gene marker clusters. Abbreviations: C, cell type; D, data set; D18, data set 18; D22A, data set 22A; N, naïve state; NL, naïve-like state(s); O, oocyte; P, primed state; T, trajectories of cellular differentiation or reprogramming, numbered as T1, T2, T2a, T2b, and T2c.

### Data transformations for meta-analysis

Meta-analysis was used to analyze the above 12 datasets in order to compare the pluripotent and differentiation states. To enable different datasets to be used for meta-analysis, we transformed all datasets by the following major steps: (i) quantile normalization within datasets using the R code, (ii) data filtering to retain protein-coding (unique) genes only, (iii) data collapse methods to calculate median values of multiple gene probes, and (iv) percentile coding (from 1 to 100%) of each gene expression in each sample.

Briefly, we performed quantile normalization (Log_2_) expression of the transcriptome per dataset (total datasets = 12) and created a master expression matrix of 16,703 protein coding genes in rows and 422 samples in columns. The expression levels recorded in this matrix by all samples have been further coded by quantile bins (1-100%). That is, the observed expression values across all genes for a sample are used to define the 1^th^ to 100^th^ quantile values. These quantile values are then used to code where each expression value for the sample falls. For example, if an expression value for a gene of the sample falls between the 20^th^ and 30^th^ quantile values, the gene expression then has a defined value of 20. This quantile-bin approach has been applied for all genes per individual sample. We next used this quantile-coded matrix to carry out meta-analysis based on the PCA, *t*-SNE, and SC3 consensus clustering, aiming to reveal the major influencing factors that control cellular and pluripotent states.

### PCA

The quantile-coded datasets were used to construct a covariance-based matrix (*A*, a gene expression versus gene expression matrix) that accounts for 100% variations of gene expression profiles. The inverse covariance matrix (*A*^*-1*^) was further employed to calculate the principal components (PCs, known as eigenvalues) (e.g., PC1, PC2, and PC3) by orthogonal decomposition using R programming. For example, PC1, PC2, and PC3 represent the sum of weighted variance (*W*_*i*_) for each individual gene expression (*Gi*) of each sample (*S*_*j*_) in one column. Thus, these PC values were used to map the data points in PCA scatter plots, in which each data point contains the genome-wide gene expression profile of one sample (or cell type). We also performed the PCA based on Pearson correlation coefficients.

### *t*-SNE visualization and cluster analysis

The *t*-SNE analysis was implemented based on a curated meta-information table, in which the details per sample can be differentiated by colored clusters in plots. For example, we can assign colors to all samples based on the names of datasets, species (e.g., mouse versus human), pluripotent states (e.g., naïve, naive-like, and primed), and cellular states (e.g., undifferentiated versus differentiated). Of note, the *t*-SNE plots may differ from each other using the same datasets for analysis at a different time (van der Maaten and Hinton, 2008).

### SC3 consensus clustering

To increase the strength of the PCA and *t*-SNE analysis, we integrated SC3 consensus clustering into PCA and *t*-SNE plots. This gene expression clustering method was based on Euclidean, Pearson, and Spearman distances using the SC3 consensus clustering (http://bioconductor.org) provided by the R package (Kiselev et al., 2017).

### Web resources used in study

https://www.ncbi.nlm.nih.gov/geo/

http://bioconductor.org/packages/release/bioc/vignettes/SC3/inst/doc/SC3.html

https://distill.pub/2016/misread-tsne/

## RESULTS

### Unsupervised PCA, *t*-SNE, and SC3 consensus clustering revealed interlaboratory data variations as a major influencing factor in assessing pluripotent and cellular states

By integrating genome-wide microarray and RNA-sequencing data into the PCA, *t*-SNE, and SC3 consensus clustering, we examined the relationships among all datasets, cell types, and cellular states in 2D plots (Figure 1, Supplemental Figures 1 and 3). One of the most intriguing findings is that the interlaboratory data variations, revealed by different principal components (e.g., PC1, PC2, and PC3), are much higher than those of intra-laboratory data with respect to cell types, pluripotent states, and gene marker clusters (Figure 1a-d). Generally, data or datasets within in a laboratory tend to group closer, regardless of the differences in their cell types.

Moreover, we observed that the paths from differentiated states toward naïve-like intermediate state *via* a major trajectory (T1). The intermediate group of cells, including primed, naïve-like, and naïve states (Figure 1c), are further projected toward naïve and naïve-like states *via* three distinct trajectories (T2a, T2b, and T2c) that may be explained by PC3 (~3.6%) (Figure 1c). To better visualize the results, we also performed *t*-SNE for the same datasets (Figure 1e-h). As a result, dataset D1, for example, can be separated from one major group (Figure 1c) into three groups, which represent one primed and two differentiation states (Figure 1g). Nonetheless, our meta-analysis suggests a significant difference between interlaboratory primed states of hPSCs (derived or maintained in different laboratories) (Figure 1).

To provide a better resolution of cellular or pluripotent states, we also employed transcriptomic cluster analysis based on expression of all unique genes (Supplemental Table 2, Supplemental Figure 3). The 30 consensus clusters (i.e., C1 to C30) enable the assignment of gene marker similarities to different pluripotent states of the cells (Figure 1d). Noticeably, closely related (e.g., C1 and C2) and distal (e.g., C17) cluster(s) tend to define a specific pluripotent state (e.g., primed) in the D1 dataset (Figure 1g). Additionally, we also generated Pearson correlation-based PCA plots (Supplemental Figure 2), which showed similar results to the above PCA analyses. Therefore, this combined approach offers a precise way to assess pluripotent states for large interlaboratory datasets that have significant systemic variations.

### SC3 consensus clustering unveiled multiple enriched small RNA clusters that define various cellular and pluripotent states

Based on the 12 datasets, we have constructed a heatmap dendrogram that is composed of the consensus clusters described above (Figure 2a-c, Supplemental Figure 3). We identified 30 individual gene clusters among these datasets after post-percentile normalization using the SC3 consensus clustering method (Kiselev et al., 2017). The dendrogram delineates the similarities or dissimilarities among 30 gene expression clusters, which are defined by the *P* values (*P* < 0.05) and the <u>a</u>rea <u>u</u>nder receiver <u>o</u>perating <u>c</u>haracteristic (AUROC) (AUROC > 0.80) (Figure 2b). The numbers of gene markers in all clusters range from 6 to 1600, with total 12,690 gene markers among the 30 clusters (Figure 2, Supplemental Table 2). To provide a quick glimpse of the potential characteristics of each cluster, we labeled the top gene marker over the box plot in each individual cluster (Figure 2b).

**Figure 2.**
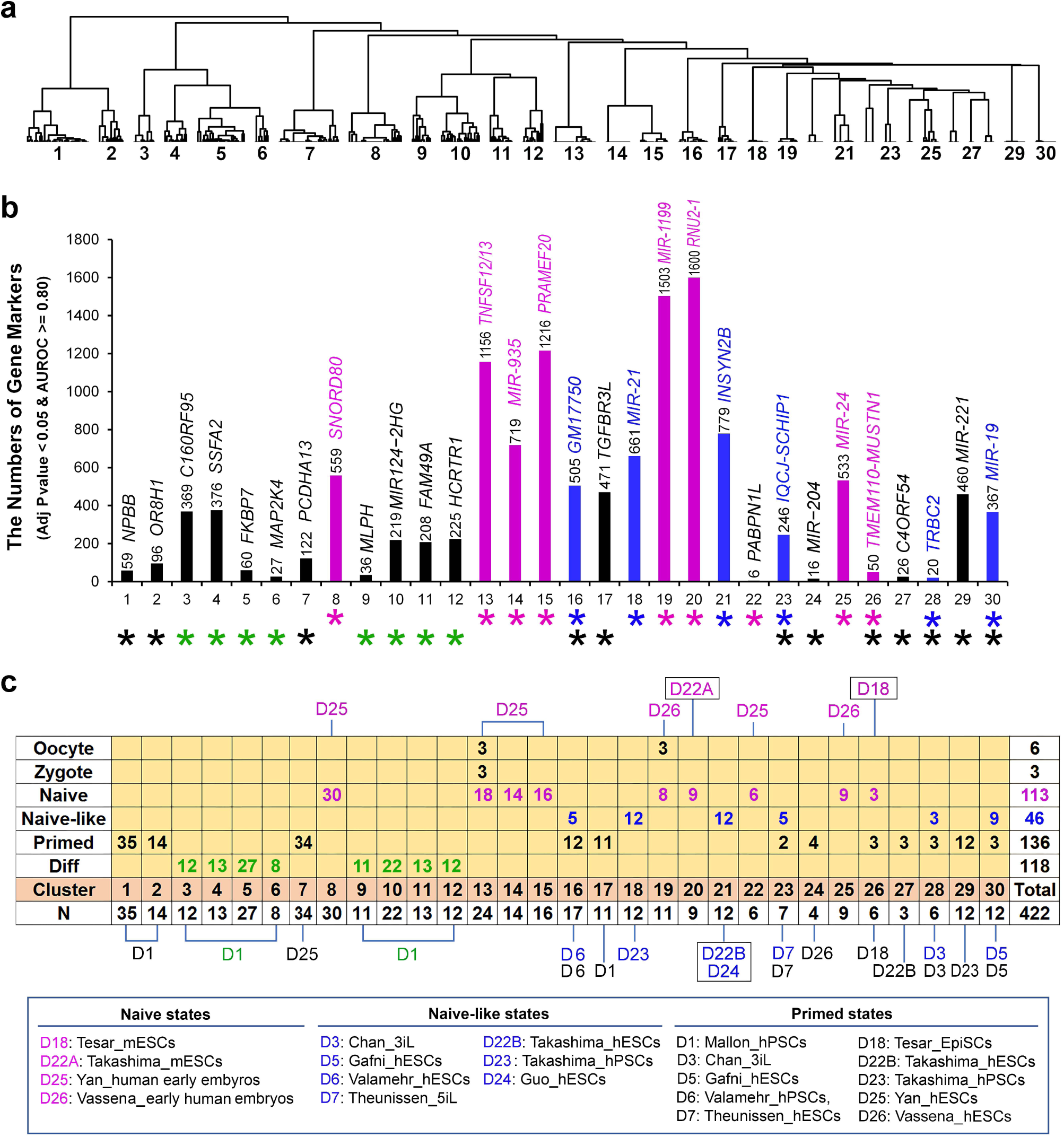
Transcriptomic clustering, gene marker identifications, and intercellular relationships among various independent studies. (**a**) Thirty individual gene clusters were among 12 datasets derived post-percentile (1-100%) datasets based on Euclidean, Pearson, and Spearman distances using the SC3 consensus clustering method (http://bioconductor.org) provided by the R package. Shown here is the dendrogram delineating the similarities among 30 gene expression clusters. The whole heatmap is available in Supplemental Figure 3. (**b**) Histogram that summarizes the numbers of gene markers in 30 clusters, which are defined by *P* values (< 0.05) and the <u>a</u>rea <u>u</u>nder receiver <u>o</u>perating <u>c</u>haracteristic (AUROC, >0.80). A top gene marker of each cluster is labeled on the top of the box plot. Purple-, blue-, and green-colored asterisk signs are used to denote the clusters defining naïve, naïve-like, and differentiated states, respectively. (**c**) Tabular presentation of the relationships between different cellular (oocyte and zygote), pluripotent (naïve and primed), and differentiated states, all of which are associated with the 30 clusters in 12 datasets (as indicated from D1 to D26). Lower panel: detailed descriptions of pluripotent states that are associated individual datasets, in which mESCs are used as naive state controls for facilitating comparative analysis. Abbreviations: Adj, adjusted; Diff, differentiation; N, the number of samples used in each cluster; Naïve, naïve pluripotent state; Naïve-like, naïve-like pluripotent state; Primed, primed pluripotent state.

Noticeably, there are 6 gene clusters, characteristic of *in vitro* cell types with naïve-like states (Figure 2b, blue asterisks), which include C16 (with top 4 gene markers depicted hereafter, ***GM17750, MIR670HG***, *UGT1A3*, and *TRAV8-1*); C18 (***MIR-21, SNORD59A, MIR-134***, and ***SNORD82***); C21 (*INSYN2B*, ***MIR-433***, *MT-TL1*, and ***MIR-668***); C23 (*IQCJ-SCHIP1, LRRC30, TRBV9*, and *PISRT1*); C28 (*TRBC2, CCDC92B, FAM243A/B*, and *SBK3*); and C30 (***MIR-19***, *HMX3*, ***MIR-218***, and *C10ORF76*) (Figure 2b, Supplemental Table 2).

Moreover, 9 gene marker clusters appear to be associated with naive states (Figure 2b, pink asterisks), which comprise C8 (***SNORD80, SNORD88C, SNORD88A***, and ***SNORD45B***), C13 (*TNFSF12/13, GLYATL3, UGT2A2*, and *TRIM6/34*); C14 (***MIR-935***, *HJV, ZNF878*, and *EIF4EBP3*); C15 (*PRAMEF20*, ***MIR-363, MIR-497***, and ***MIR-675***); C19 (***MIR-1199***, *C4ORF50*, ***MIR-124***, *and FAM227A*); C20 (***RNU2-1, SNORA43, SNORA17***, and ***MIR-290***); C22 (*PABPN1L, ZNF705A, UBTFL1*, and *OTULINL*); C25 (***MIR-24***, *TEX52, TMEM247*, and ***MIR-149***), and C26 (*TMEM110-MUSTN1, ISY1-RAB43, MFSD13B*, and *TRGV2*) (Figure 2b, Supplemental Table 2).

Interestingly, among the above-mentioned 15 gene clusters that classify naive and naïve-like states, there are 67% (n = 10) which have RNA and/or microRNA markers within the top 4 gene marker list (see above bold gene symbols and Supplemental Table 2). Particularly, C8, C15, C18, and C20 are enriched with small nucleolar RNAs (snoRNAs) (Supplemental Table 2), which belong to noncoding RNAs involved in the processing and modification of ribosomal RNAs and microRNAs. These data indicated that a large amount of RNA and small RNA markers may be used as novel markers to characterize the naïve and naive-like states.

To reveal the complicated relationship between different cellular (e.g., oocyte and zygote), pluripotent (e.g., naïve, naïve-like, and primed), and differentiated states, we linked cellular state data to both SC3 consensus clusters and the cellular identities from the 12 datasets for a comparative analysis. As shown in Figure 2c, all cellular and pluripotent states can be assigned into all 30 clusters. Convincingly, two differentiation groups in D1 are individually grouped by two consecutive clusters (i.e., C3-6, C9-12). Furthermore, this comparative analysis confirms the significant heterogeneity among naïve, naïve-like, and primed states. For examples, naïve states are associated with the clusters spanning from C8 to C26, which are adjacent to D25 (processed by the RNA-sequencing platform) and D26 (processed by cDNA microarray analysis) respectively (Figure 2c). Thus, the cluster-dataset relationship may be explained by the use of different laboratory protocols used for analyzing RNA expression.

Markedly, naïve mESCs from both D18 and D22A also clustered significantly differently, with a distance of 5 clusters. Moreover, naïve and naïve-like state clusters do not overlap, suggesting that hPSCs with naïve and naïve-like states have different pluripotent states. Within the datasets D3, D5, D6, and D7, cell samples with naïve-like states share some clusters with primed states, suggesting that these naïve-like hPSCs have a closer relationship with primed hPSCs than naïve hPSCs. Interestingly, C17 distinguishes 11 primed hESC lines from their hiPSC counterparts (Figure 2c), with significant differences between 471 gene markers (as topped by *TGFBR3L, SHISA8, IGLC1*, and *SDCCAG3*) (Supplemental Table 2).

### Top gene markers show relative specificity for discerning pluripotent and cellular states

To evaluate the relative specificity of top gene markers among the 30 clusters, we calculated the mean standardized expression (ranging 1-100%) of each gene marker in all samples under each cluster (Figure 3). It appears that the top gene marker expression in the first 13 clusters (i.e., C1-C15, except C8 and C15) lacks specificity for a particular gene cluster, displaying a significant fluctuation/scatter among the 30 clusters. However, the expression of top gene markers from C15 to C30 exhibits much higher specificity demonstrated by lower scatter (Figure 3).

**Figure 3.**
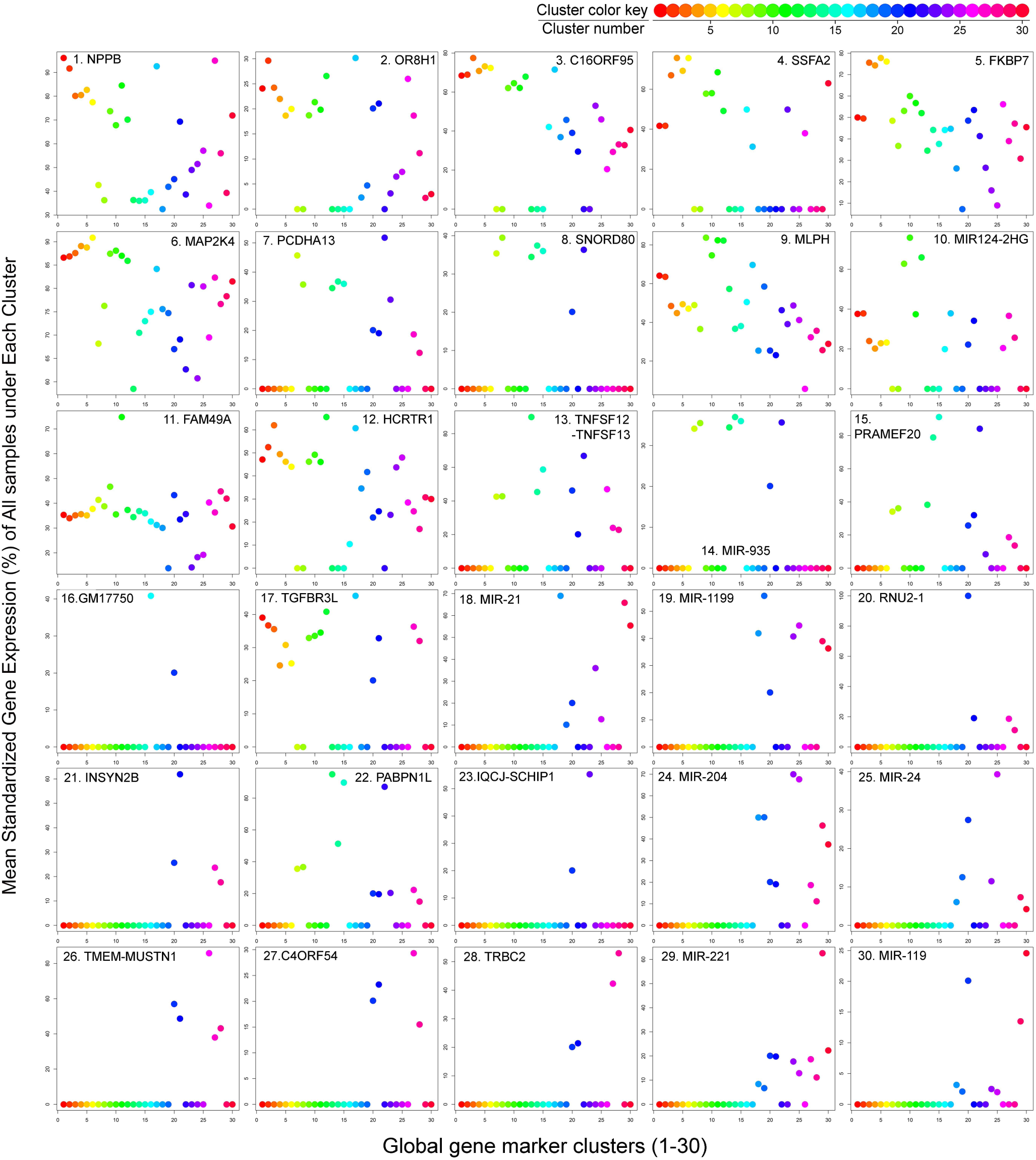
Top gene marker presentation in the 30 unsupervised individual clusters. Each color dot represents mean standardized gene expression (1-100%) of all samples per cluster. Only one top marker is labeled in the plot.

This increased specificity seems to be associated with the expression of RNA or microRNA genes (e.g., *GM17750, MIR-1221, MIR-1199, RNU2-1, MIR-204, MIR-24, MIR-221, and MIR-119*) in naïve, naive-like, and primed states (Figure 3). For example, *GM17750* is specific to C16 (which connected to Valamehr naïve-like and primed hESCs, D6), *RNU2-1* to C20 (Takashima mESCs, D22A), *MIR-221* to C29 (Takashima primed hPSCs, D23), and *MIR-119* to C30 (Gafni naïve-like and primed hPSCs, D5) (Figures 2c and 3). Thus, the relative specificity of top gene marker expression would enable its utility to define cellular identities, states, and potential functions in NLPs that are generated from different laboratories.

### Supervised cluster analysis found new gene signatures for characterizing pluripotent and cellular states

Beside the above unsupervised meta-analysis, we also present here 4 supervised heatmaps that delineate normalized mean expression of 88 known genes (www.genecards.org), which encode dominant signaling molecules (n = 26, Figure 4a), differentiation markers (n = 25, Figure 4b), developmental regulators (n = 25, Figure 4c), and key pluripotent transcriptional factors (n = 12, Figure 4d). These heatmaps demonstrate new features of cellular and pluripotent similarities associated with the 30 gene clusters as described in Figure 2a.

**Figure 4.**
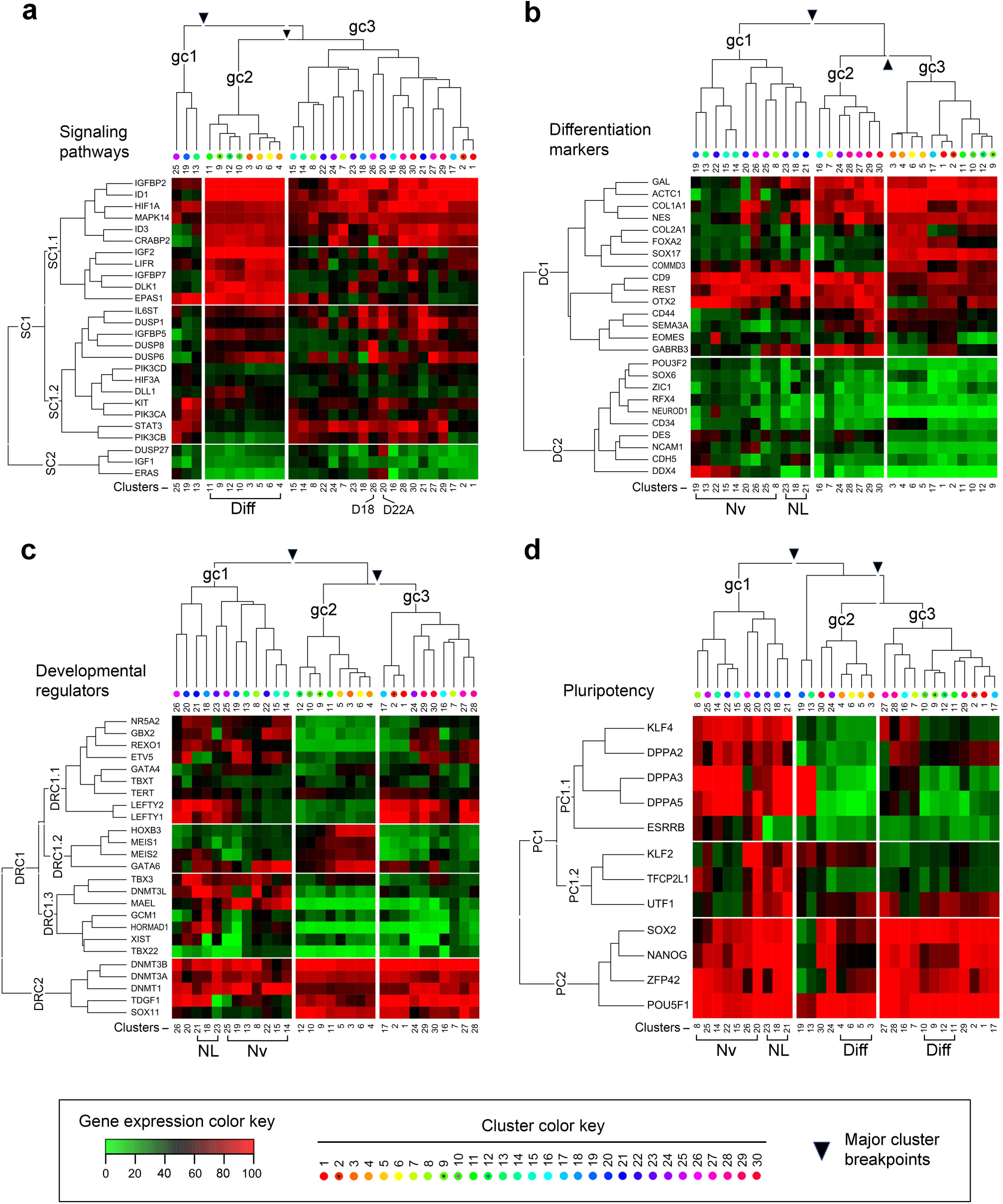
Heatmap of normalized mean expression for supervised gene markers for all samples under each defined cluster. Shown in the right panel is a heatmap of normalized mean expression (1-100%) across all clusters for supervised gene markers. Also see Supplemental Table 2 for details. Abbreviations: D18, data set 18; D22A, data set 22A; DC, differentiation gene cluster; Diff, differentiation or differentiated state(s); DRC, developmental regulator gene cluster; gc, global gene cluster; N, naïve pluripotent state; NL, naïve-like pluripotent state; PC, Pluripotency gene cluster; SC, signaling pathway cluster.

We initially analyzed the influence of gene markers that encode common signaling pathways. As indicated by annotated major cluster breakpoints, mean standardized expression of these markers results in three major signaling gene clusters (SC1.1, SC1.2, and SC2) that classify the 30 consensus clusters, derived from the unsupervised PCA and *t*-SNE, into three global clusters (gc1, gc2, and gc3) that define differentiation and pluripotent states (Figure 4a). The gc1 contains C13, C19, and C25, defining a unique human naïve state shared by both Vassena and Yan datasets (Vassena et al., 2011; Yan et al., 2013). The human naïve (C13, C19, and C25), differentiation (C3-6 and C9-12), and naïve-like clusters appear to be driven by expression of a 11-gene cluster (*IGFBP2, ID1, HIF1A, MAPK14, ID3, CRABP2, IGF2, LIFR, IGFBP7, DLK1*, and *EPAS1*) in the SC1.1 block (Figure 4a). Interestingly, upregulation of the 5-gene subset (*IGF2, LIFR, IGFBP7, DLK1*, and *EPAS1*) is a strong indicator for two differentiation states in D1. However, these signaling gene markers do not provide clear signatures to distinguish naïve from naive-like and primed states.

We next examined the impact of differentiation gene markers on the rearrangements of the 30 clusters. As shown by the cluster breakpoints, mean standardized expression of these gene markers results in two major differentiation clusters (DC1 and DC2) that organize the 30 consensus clusters into three global clusters (i.e., gc1, gc2, and gc3) (Figure 4b). Under this analysis, both DC1 and DC2 are able to cluster all naïve states as well as naïve-like states into gc1 (Figure 4b). Moreover, the two clusters C20 and C26, which contain the two mESC datasets, bear most resemblance to each other under this supervised condition, which is in contrast to our unsupervised analyses.

Concerning the influence of developmental regulators on the relocations of the 30 clusters, three major DNA methyltransferase genes (i.e., *DNMT1, DNMT3A*, and *DNMT3B*) showed ubiquitously high levels of mRNA expression in these 30 clusters (Figure 4c), thus diminishing their predictive values for the pluripotent states. However, the predictive values for both naïve and naïve-like states might underlie the expression pattern of *TBX3, DNMT3L*, and *MAEL* within the DRC1.3 cluster (Figure 4c). Evidently, high level of *Xist* transcripts, one of the hallmarks of naïve pluripotency, is only found in both C20 and C21, which defines the datasets containing both naïve mESCs and primed EpiSCs (Figure 4c).

Finally, we interrogated the effect of known pluripotent regulators on gene cluster rearrangements. We found that three well-established pluripotent regulators (i.e., *SOX2, NANOG, and POUF51*), clustered with a poorly-understood regulator *ZFP42*, had no predicative values for discriminating diverse pluripotent states (Figure 4c). However, the predictive values for both naïve and naïve-like states clearly underlie the expression pattern of the PC1.1 (*KLF4, DPPA2, DPPA3, DPPA5*, and *ESSRB*) and PC1.2 (*KLF2, TFCP2L1*, and *UTF1*) clusters (Figure 4c). In particular, *ESRRB* expression patterns distinguish naïve pluripotent states from those naïve-like, primed, and differentiation states. Moreover, the 3-gene cluster *KLF2, TFCP2L1*, and *UTF1* discriminates mESCs (C20 and C26) from human naïve pluripotency, which is defined by datasets from early human embryos (see C8, C14, C15, C22, and C25). Thus, these supervised analyses not only confirm the value of some previously identified gene markers (such as *KLF2, TFCP2L1*, and *ESSRB*) but also unveil new gene signatures or combinations for discerning pluripotent states.

## DISCUSSION

Achieving human naive pluripotency through the perturbation of growth factor signaling is believed to have significant impacts on hPSC growth, expansion, genetic engineering, disease modeling, and drug discovery. However, to accurately define a pluripotent state seems to be hindered by a lack of reliable analytic tools for comparative meta-analysis. Based on the integrated meta-analysis described in this study, we have provided an unbiased evaluation of the rationale and reliability of using meta-analysis for the assessment of cellular and pluripotent states. Here, we will discuss critical influencing factors, integration of supervised into unsupervised analyses, and several new findings based on this meta-analysis.

### The rationale and reliability of using meta-analysis for the assessment of cellular and pluripotent states

To enable impartial and meaningful comparison of the transcriptomic levels of interlaboratory datasets, a pre-processing strategy must first be employed. Here, we demonstrate one such pre-processing strategy that involves quantile normalization of mRNA expression across samples within each dataset followed by percentile coding of the normalized gene expression. By doing so, the ranking of mRNA expression within a sample for a dataset is preserved and the resulting expression transformed to a value between 1 and 100. These transformed values can then be used to directly compare ranks of mRNA expression across datasets without a substantial bias. Moreover, the post-transformed values for different datasets generated from different laboratories using different technologies can be collectively visualized using both PCA and *t*-SNE, which are subjected to unsupervised analysis using SC3 consensus clustering.

The differences between primed and differentiated states (within and between datasets) remain preserved after such pre-processing and, for some datasets, each state can be further subclustered (Figures 1–5). Moreover, the transformed values for each mRNA can be reanalyzed and representative markers for each cluster identified, including those that discriminate the pluripotent state (e.g., primed and naïve) and source of cells (e.g., hESC and hiPSC). The reliability of this analysis was demonstrated in PCA, *t-*SNE, and SC3 consensus clustering in a large control dataset (D1) from the NIH Stem Cell Unit, in which both primed and differentiated states were clearly separated with various relevant clusters (Figures 1–5). Likewise, the differences between hiPSC and some hESC lines may be revealed by gene marker expression in cluster 17 (C17) (Figure 2c), which possesses *TGFBR3L, SHISA8, IGLC1*, and *SDCCAG3* as top gene markers, a novel finding that has not been not previously reported.

### Integration of supervised into unsupervised analyses: *pros* and *cons*

To better view the relationship between clustering analysis and pluripotent states, we integrated the supervised into unsupervised analyses by introducing cluster breakpoints and by aligning the clusters with cellular or pluripotent state information (Figure 5). The cluster fragments provide a quick view of the interchangeability of cluster rearrangements. As indicated in Figure 5, the rearrangements of gene clusters depend on the size(s), function, and redundancy of input gene clusters. Combining SC3 consensus clustering with both PCA and *t*-SNE, we may integrate supervised into unsupervised analyses in a one-dimensional format (Figure 5). Thus, this dimensional reduction provides a rapid, informative, and unbiased view of cellular and pluripotent states under various interlaboratory growth and assay conditions.

**Figure 5.**
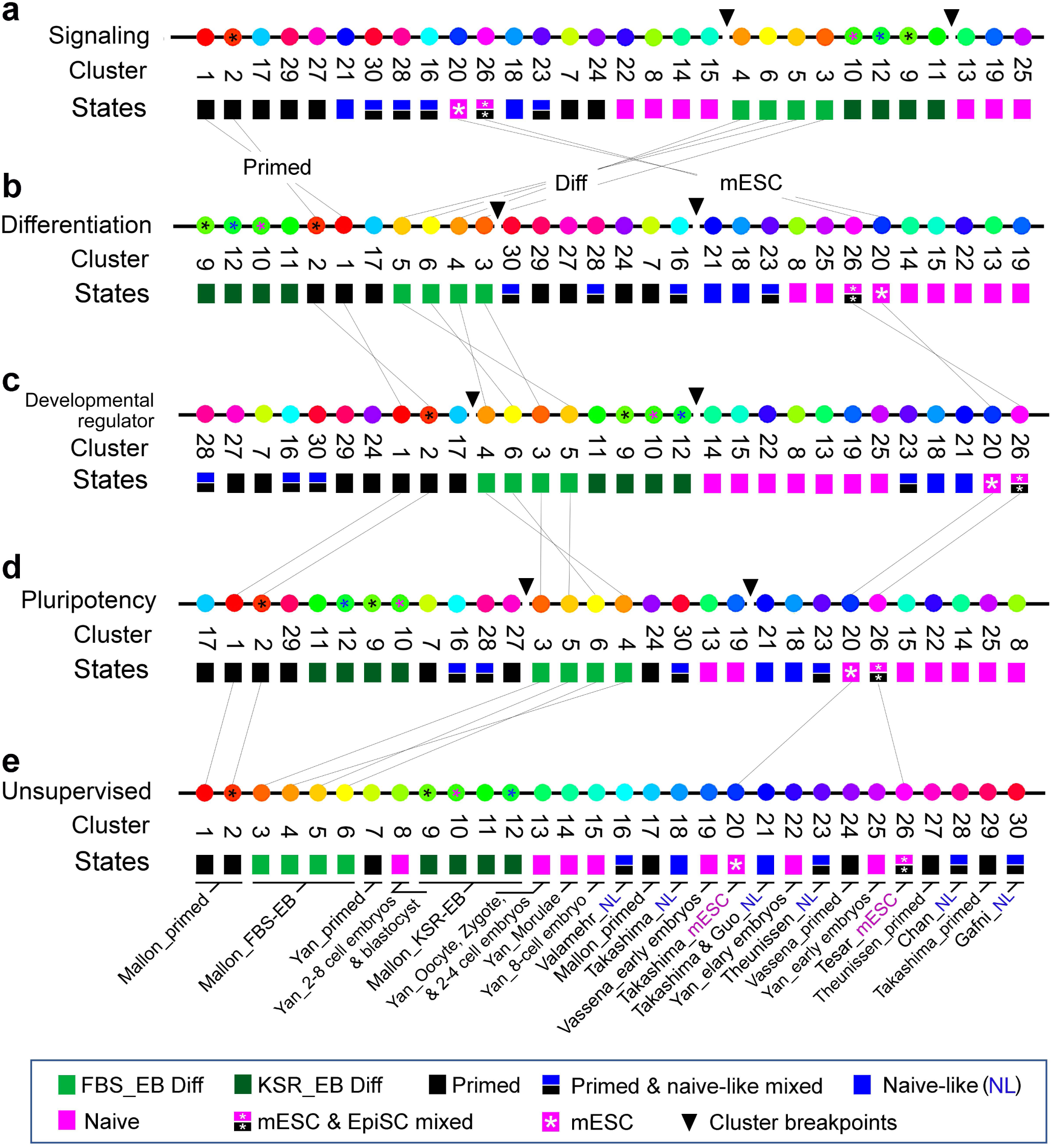
One-dimensional mapping of pluripotent and cellular identities by integrating cellular states with cluster analysis from independent studies. Black arrowheads indicate cluster breakpoints that enable cluster rearrangements. Abbreviations: EpiSC, primed mouse epiblast stem cells; FBS_EB Diff, fetal bovine serum (FBS)-mediated embryoid body (EB) differentiation; KSR_EB Diff, KnockOut™ Serum Replacement (KSR)-mediated embryoid body (EB) differentiation; mESC, naïve mouse embryonic stem cells; Naïve, naïve pluripotent state; NL, naïve-like pluripotent state; Primed, primed pluripotent state.

For example, mESC naïve cells (from D18) showed the closest relationship (as indicated by C26) with primed EpiSC control from the same dataset in clustering analysis (Figure 2a and Supplemental Figure 3). However, the two naïve mESC controls (from D18 and D22A) do not show a close relationship in either PCA plots or SC3 consensus clustering (Figures 1d, 1h, and 2). This discrepancy between the two mESC lines may be explained by their different cell culture methods, in which the mESCs from D18 were cultivated under a normoxic condition (19% O_2_) whereas the mESCs from D22A were maintained in hypoxia (i.e., 5% O_2_) (Tesar et al., 2014; Takashima et al., 2014).

Moreover, in another independent meta-analysis, Smith and colleagues showed human naïve-like Reset cells (induced by 2iL and the PKC inhibitor Gö6893) were similar to their naïve mESCs in PCA plots (Takashima et al., 2014). In their meta-analysis, they indicated that NLPs (from both Gafni and Chan datasets), located distal from the Takashima datasets in the PCA plot, are more similar to their primed counterparts (Takashima et al., 2014). Clearly, our meta-analysis revealed much larger differences between interlaboratory datasets than their actual cellular state differences between intra-laboratory cell samples, suggesting the possibility that the previously reported similarities or differences in naïve and naïve-like cellular models are likely attributed to interlaboratory protocol differences.

Importantly, the number of gene markers used for conducting unsupervised or supervised clustering analysis has a significant impact on the relationship among various cellular and pluripotent states. A statement made from ether a supervised (usually with a small subset of markers) or unsupervised analysis (e.g., in this study) should be weighed differently. However, we recommend integrating the supervised into unsupervised analyses to accurately interpret interlaboratory datasets (Figure 5).

### Critical influencing factors of meta-analysis in the stem-cell field

Beside the interlaboratory protocol differences that may derail a successful meta-analysis, several other influencing factors should also be taken into consideration when implementing meta-analysis. These factors include experimental data variability attributed to data types (e.g. cDNA microarray and RNA-seq), interlaboratory data standardization and normalization, data stabilization, and the choices of methods (e.g., PCA or *t*-SNE) used for meta-analysis.

For example, when dealing with cDNA microarray and RNA-sequencing data for the analysis, we frequently encounter: (i) sensitivity related to Poly A in channel versus ribosomal RNA depletion; (ii) sequencing depth (e.g., coverage, numbers of reads, and the length of reads used for mapping); (ii) paired versus single ends (in which paired ends are more accurate); (iv) stranded versus non-stranded; (v) and outliers. Accordingly, the distinct SC3 consensus clustering differences (C13-15 versus C25) between the early human embryo datasets D25 (RNA-sequencing) and D26 (cDNA microarray) seem to be consequential to one of the above discussed issues (Figure 2). Thus, more reliable data normalization and/or transformation methods should be developed to overcome these technical problems prior to its broad applications in the stem-cell field.

With respect to the PCA, separation of interlaboratory data in the plots seems to be insufficient when including multi-laboratory datasets for the analysis. This problem may be overcome by comparing with the *t*-SNE and SC3 consensus clustering (Figure 1). Concerning *t-*SNE visualization and cluster analysis, the high-dimensional data reduction technique, originally developed by van der Maaten and Hinton in 2008 (van der Maaten and Hinton, 2008), has gained popularity for data analysis and machine learning in recent years. The greatest advantage of the *t*-SNE lies in its ability to visualize data in a fascinating 2D plot for high-dimensional datasets (up to thousands of dimensions). However, we should be aware that this technique is a random and non-linear method. Its dimensions, physical distances of data points, size of clusters are meaningless in the plots (https://distill.pub/2016/misread-tsne/). Noticeably, *t*-SNE plots may differ from each other in the same datasets. To enhance the strength of *t*-SNE in meta-analysis, we emphasize the combined use of *t-*SNE with both PCA and SC3 consensus clustering as demonstrated in this study.

## CONCLUDING REMARKS

Our meta-analysis indicates that hPSCs grown under current naïve-like protocols do not have naïve pluripotent clusters as described by mESCs and early human embryos. This may be due to the considerable heterogeneity among various cellular and pluripotent states in a large cohort of datasets generated from different laboratories. Thus, interlaboratory data variation represents the predominant factor interfering with the accuracy of meta-analysis, which significantly limits the predictive values of meta-analysis for defining cellular and pluripotent states.

The combined use of percentile normalization with PCA, *t*-SNE, and SC3 consensus clustering, which represents one new strategy to compare multiple interlaboratory data sets. has significantly improved the predictive values of the current meta-analysis. Other data normalization or transformation algorithms (e.g., using highly consistent housekeep genes in each cell type prior to the percentile normalization as described in this study) should be considered in the future, which would be crucial for reducing interlaboratory data disparities. No doubt, standardization of interlaboratory protocols for stem cells and development of new data normalization algorithms would enable a robust and reliable meta-analysis of cellular and pluripotent states.

## ACKNOWLEDGMENTS

This work was supported by the Intramural Research Program of the NIH at the National Institute of Neurological Disorders and Stroke. We thank Dr. Kyeyoon Park, Dr. Paul Tesar, and Dr. Pamela Robey for helpful discussions.

---

**Supplemental Figure 1.**
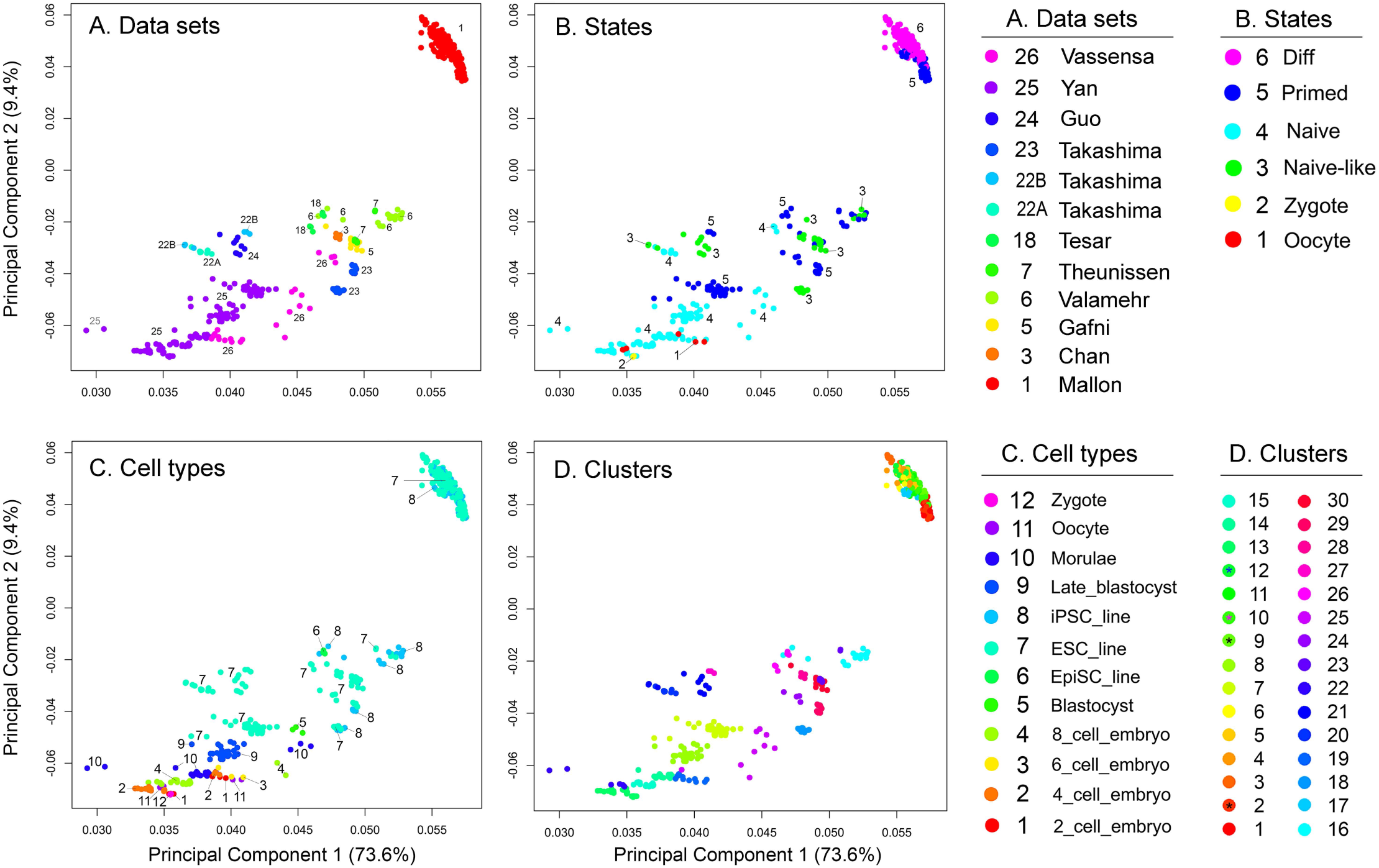

**Supplemental Figure 2.**
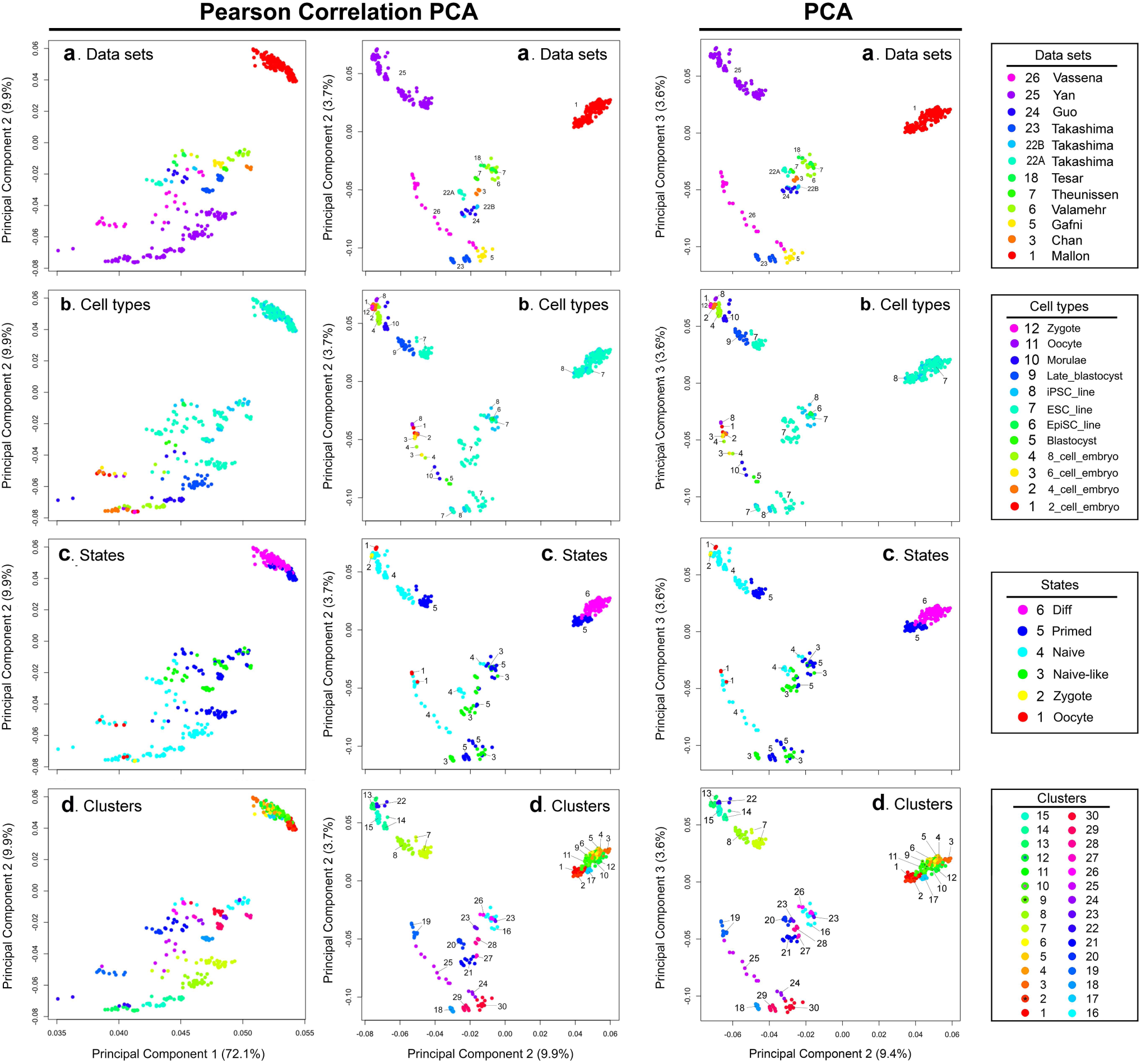

**Supplemental Figure 3.**
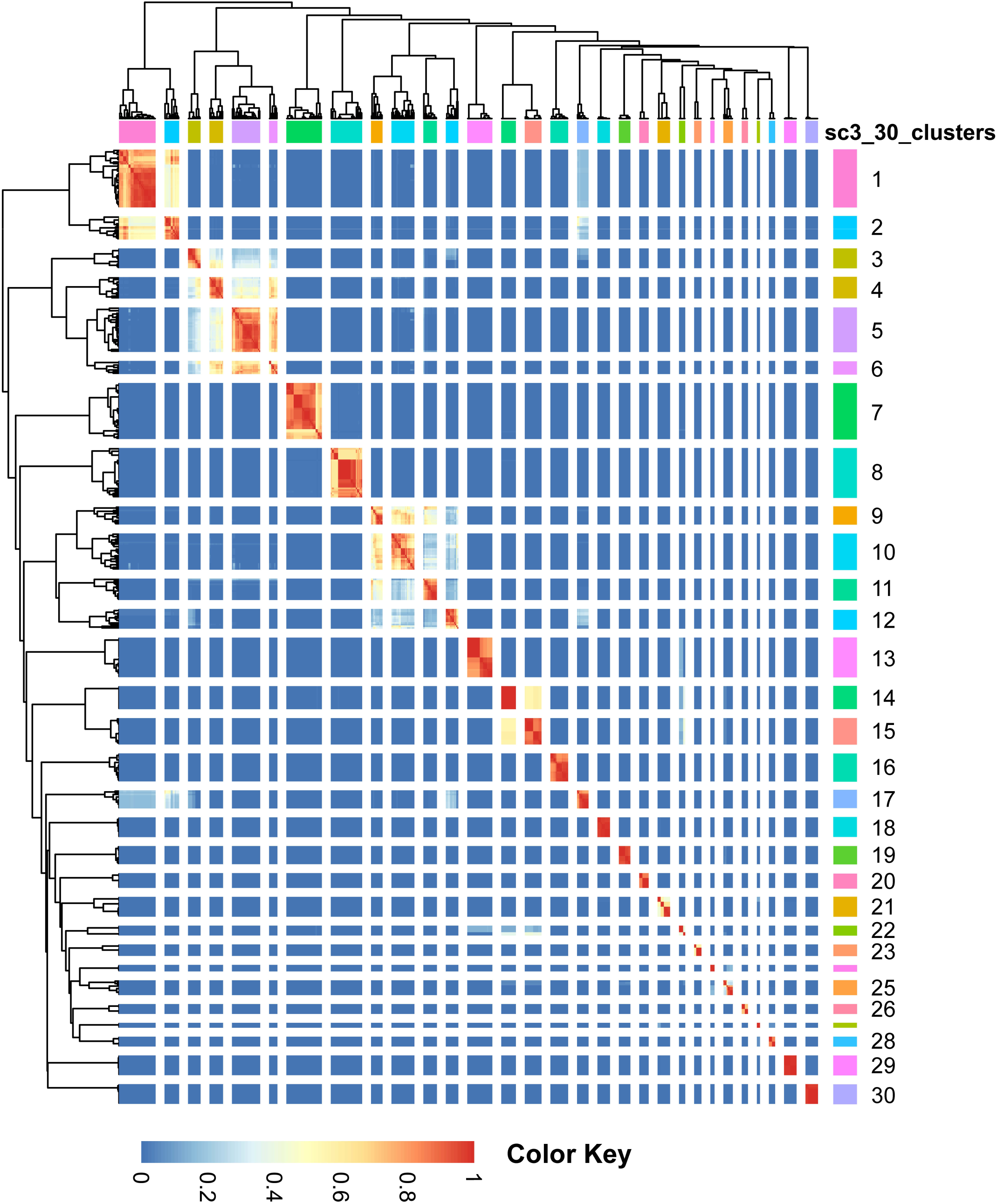

